# Hybridization preceded radiation in diploid wheats

**DOI:** 10.1101/599068

**Authors:** Stella Huynh, Thomas Marcussen, François Felber, Christian Parisod

## Abstract

Evolutionary relationships among the *Aegilops-Triticum* relatives of cultivated wheats have been difficult to resolve owing to incomplete lineage sorting and reticulate evolution. Recent studies have suggested that the wheat D-genome lineage (progenitor of *Ae. tauschii*) originated through homoploid hybridization between the A-genome lineage (progenitor of *Triticum* s.str.) and the B-genome lineage (progenitor of *Ae. speltoides*). Scenarios of reticulation have been debated, calling for adequate phylogenetic analyses based on comprehensive sampling. To reconstruct the evolution of *Aegilops-Triticum* diploids, we here combined high-throughput sequencing of 38 nuclear low-copy loci of multiple accessions of all 13 species with inferences of the species phylogeny using the full-parameterized MCMC_SEQ method. Phylogenies recovered a monophyletic *Aegilops-Triticum* lineage that began diversifying ~6.5 Ma ago and gave rise to four sublineages, i.e. the A- (2 species), B- (1 species), D- (9 species) and T- (*Ae. mutica*) genome lineage. Full-parameterized phylogenies as well as patterns of tree dilation and tree compression supported a hybrid origin of the D-genome lineage from A and B ~4.1 Ma ago, and did not indicate additional hybridization events. This comprehensive and dated phylogeny of wheat relatives indicates that the origin of the hybrid D-genome was followed by intense diversification into almost all diploid as well as allopolyploid wild wheats.

## 1. Introduction

Triticeae is one of the most economically important tribes of the grass family, Poaceae, and includes such globally significant cereal crops as wheats (*Triticum* spp.), rye (*Secale cereale*) and barley (*Hordeum vulgare*). Next to cultivated wheats, wild wheat species comprise a dozen of annuals from the genera *Aegilops* and *Triticum* that occur naturally across the Mediterranean region and into Central Asia (Kilian et al., 2011; van Slageren, 1994). Efforts to analyze the genomes of cultivated and wild wheats to uncover their evolutionary relationships have been on for decades (Eig, 1929a; Kihara, 1954). These research efforts have largely been driven by the key role of wild wheats as potential sources for improvement of the existing cultivars. Hence, special attention has been attributed to hexaploid bread wheat (*T. aestivum* with BBAADD genome composition) and its wild diploid progenitors, i.e. *Ae. tauschii* (DD), *T. urartu* (AA) and the donor of the B subgenome which may have been closely related to *Ae. speltoides* (SS) (e.g. Bernhardt et al., 2017; Daud and Gustafson, 1996; Marcussen et al., 2014; Petersen et al., 2006).

In spite of repeated efforts, the phylogenetic relationships among diploid wheats have not yet been resolved. This should probably be seen in light of recent diversification, high coalescence stochasticity among unlinked genes, and past and present hybridization and introgression (Escobar et al., 2011; Parisod et al., 2013; Senerchia et al., 2016; Zohary and Feldman, 1962). In a phylogenetic analysis of 275 low-copy nuclear loci for hexaploid *T. aestivum* and five diploids, Marcussen et al. (2014) recovered three principal genome lineages, which they referred to as the A-, B- and D-genome lineages based on the nomenclature of wheat subgenomes. The A-genome lineage comprises *T. monococcum, T. urartu* and the *T. aestivum* A-genome, the B-genome lineage *Ae. speltoides* and the *T. aestivum* B-genome, and the D-genome lineage *Ae. sharonensis, Ae. tauschii* and the *T. aestivum* D-genome. Tree topology ratios and speciation times indicated that the D-genome lineage had originated by hybridization between the other two lineages. This study also presented the first species-level phylogeny for all diploid wheats, using a multispecies coalescent (MSC) analysis on six loci. Topological conflicts between this 6-gene phylogeny, the 275-gene phylogeny and the chloroplast phylogeny were subsequently discussed and interpreted as evidence of additional hybridization events in the history of the wheat lineage (Li et al., 2015a, 2015b; Sandve et al., 2015). El Baidouri et al. (2017) reached a similar conclusion based on transposable elements. Additional hybridizations have however not yet been subject to rigorous phylogenetic testing, as such analyses require a comprehensive sampling of taxa and a high number of genes.

Recent years have brought theoretical advances and also new methods and techniques for detecting ancient hybridization events and reconstructing phylogenetic networks from gene data in the presence of incomplete lineage sorting (see Elworth et al., 2018). These include full-parametrised, Bayesian approaches that employ the multispecies network coalescent (MSNC) to generate species networks from multilocus sequences (Wen and Nakhleh, 2018; Zhang et al., 2018). These methods are however computationally intensive and therefore numerous simplified approaches have been privileged so far. The most popular approaches take either gene trees as input and estimate a species network from tree topology ratios by maximum parsimony, maximum likelihood or Bayesian inference (e.g. Solís-Lemus et al., 2017; Yu et al., 2014, 2013), or suggest hybridization within data subsets based on SNPs (Durand et al., 2011; Green et al., 2010; Pease and Hahn, 2015). The lack of an explicit time scale can however make these methods susceptible on inherent model violations.

Here we took advantage of recent methodological advances to revisit the phylogenetic relationships among wheats and relatives. We thus developed an amplicon sequencing approach to accurately genotype 38 single-copy nuclear loci among all 13 diploid species of the *Aegilops-Triticum* wheat lineage and four outgroups, and present a dataset matching the specificities of full-parametrised inferences of relationships. Species networks inferred under the MSNC model using the MCMC_SEQ method (Wen and Nakhleh, 2018) clearly identified reticulation at the origin of the largest lineage of *Aegilops* species (the D-genome lineage including the bread wheat progenitor *Ae. tauschii*).

## 2. Material and methods

### 2.1 Plant material

A total of 19 accessions of 17 species of *Aegilops* L. and *Triticum* L. and four nested outgroups were obtained from the ARS-GRIN and ICARDA germplasm databases and used for analysis (Supplementary Table S1). Ingroup species were *Ae. bicornis* Jaub. & Spach (genome S^b^), *Ae. caudata* L. (C), *Ae. comosa* Sm. (= *Ae. markgrafii* (Greuter) K. Hammer) (M), *Ae. longissima* Schweinf. & Muschl. (S^l^), *Ae. mutica* Boiss. (= *Amblyopyrum muticum* (Boiss.) Eig) (T), *Ae. searsii* Feldman & Kislev (S^s^), *Ae. sharonensis* Eig. (S^l^), *Ae. speltoides* Tausch (S), *Ae. tauschii* Coss. (D), *Ae. uniaristata* Vis. (N), *Ae. umbellulata* Zhuk. (U), *T. monococcum* L. subsp. *monococcum* and subsp. *aegilopoides* (Link) Thell. (= *T. boeoticum* Boiss.) (A^m^), and *T. urartu* Thumanjan ex Gandilyan (A^u^). Outgroups were *Taeniatherum caput-medusae* (L.) Nevski, *Agropyron mongolicum* Keng, *Secale cereale* L. and *Hordeum vulgare* L. Species and genomic designations follow Barkworth and von Bothmer (2009). Plants were all grown outdoor at the Botanical garden of Neuchâtel until flowering and deposited as herbarium sheets at the Botanical Museum and Garden of Lausanne, Switzerland (LAU). Leaves from young seedlings were sampled and dried in silica gel for DNA extraction using the DNAeasy kit from Qiagen^®^ following the manufacturer protocol. DNA was quantified using Nanodrop and normalized at 50 ng/μL.

Up to six accessions were selected for each of eleven diploid *Aegilops* L. species and two diploid *Triticum* L. species as well as the four outgroups (Supplementary Table S1). Those 38 accessions were obtained from ARS-GRIN or ICARDA germplasm databases and were all grown outdoor at the Botanical garden of Neuchâtel until flowering and deposited as herbarium sheets at the Botanical Museum and Garden of Lausanne, Switzerland (LAU). Leaves from young seedlings were sampled and dried in silica gel for DNA extraction using the DNAeasy kit from Qiagen^®^ following the manufacturer protocol. DNA was quantified using Nanodrop and normalized at 50ng/μL.

### 2.2 Amplicon sequencing of nuclear markers

A total of 48 candidate low-copy nuclear loci were selected (Supplementary Table S2) based on available evidence from the literature (Duarte et al., 2010; Sang, 2002). For each locus, sequences from *Aegilops tauschii* (DD) and *Triticum aestivum* (BBAADD) were retrieved from assembled genomes available on EnsemblPlants (Kersey et al., 2014) through either their annotation or the best match following blastN using the corresponding locus from *Triticum aestivum*. Available sequences for each locus were aligned using MAFFT (Katoh, 2002) and primers were designed in exons encompassing 400 bp of mostly intronic regions using Primer3 (Untergasser et al., 2012). Primer pairs were designed to similarly achieve PCR amplification at a T_m_ around 60°C and were tagged with a 22-bp sequence designed by Fluidigm for the Access Array barcode library.

Simultaneous PCR amplifications of all 48 loci were performed on all samples in the 48.48 Access Array Integrative Fluidic Circuit (IFC) chip following Fluidigm instructions. Two replicates (i.e. 5.1 % of the whole dataset) and one negative control were included. The IFC chip mixed the samples and primer pairs in all possible combinations for amplification into independent nanochambers before pooling resulting amplicons for each sample. Each sample pool was barcoded with a specific 10 bp-sequence attached at one side of the amplicons through PCR (7 cycles) and cleaned using AMPure beads (Agilent) at x0.7 ratio. Quality was checked using High-Sensitive chips on the BioAnalyzer 2100 (Agilent) and sample pools were quantified through real-time PCRs using the LightCycler System in 384-well plates with four standards. All samples were finally pooled at equimolar concentrations by individual pipetting, cleaned with AMPure beads and checked for quality with BioAnalyzer 2100. The concentration of the final pool was estimated through qPCR with four replicates and normalized at 2 nmol before amplicons were sequenced using the MiSeq Reagent Kit v3 for paired-ends reads sequencing (2 x 300 bp), with an expected 200 bp overlap over the 400 bp amplicons. Following Fluidigm recommendations, custom Locked Nucleic Acid (LNA™) oligonucleotides from Exiqon were used at 10 μM to enhance specificity and sensitivity of the sequencing and PhiX Control v3 sequencing library from Illumina was used to promote quality.

### 2.3 Genotyping of nuclear markers

Overlapping paired-end reads were merged and further filtered with a mean Quality Control of 20 using BBmerge script v8.8 in BBTools (http://jgi.doe.gov/data-and-tools/bbtools/). After demultiplexing per sample (based on barcodes without mismatch) using BBmap v.35.43 in BBTools, merged reads were retrieved at each locus (based on primers with up to 1 mismatch) using CutAdapt v.1.5 (Martin, 2011) and locally aligned with Muscle (Edgar, 2004). Alignments were additionally cleaned with a custom R script (Supplementary Material Script S1) based on frequencies of nucleotides, gaps and unknown base at each position. Variants present in less than 10% of the reads were removed and remaining reads were characterized into haplotypes for each locus. Individuals presenting two haplotypes at a given locus were considered heterozygous.

Corresponding sequences of the distant Pooideae outgroup *Brachypodium distachyon* were retrieved from the available reference genome (v1.0 INSDC assembly; The International Brachypodium Initiative, 2010) and added to our dataset. They were then iteratively aligned for each locus using SATé (Liu et al., 2009) with the following settings: MAFFT as Aligner, Muscle as Merger, RAxML (Stamatakis, 2014) as Tree Estimator, GTRGAMMAI as nucleotide substitution model, and SATé-II-ML with Centroid decomposition and “best alignment after last improvement”. After indel adjustments using AliView v1.25 (Larsson, 2014), the final alignments had a total aligned length of 18046 bp.

### 2.4 Phylogenetic analyses

A combination of phylogenetic approaches and analyses were used to identify and test for hybridization in the wheat lineage: full-parameterized analyses using MCMC_SEQ (Wen and Nakhleh, 2018), analysis of gene tree topology ratios using SNaQ (Solís-Lemus et al., 2017), and analysis of SNP ratios using ABBA BABA-tests (Durand et al., 2011; Green et al., 2010).

#### 2.4.1 Global visualization of sequence data

We first generated an exploratory neighbor-net splits network (Bryant and Moulton, 2002) in SplitsTree4 (Huson and Bryant, 2006), based on SNP differences (i.e. uncorrected P) between concatenated genes for all individuals without phasing. We deemed one random allele per locus from each of the 19 taxa in Table S1 and *Brachypodium distachyon* as sufficient and adequate sampling for the downstream, computationally demanding analyses.

#### 2.4.2 Full-parameterized inference of phylogenetic network

The MCMC_SEQ method (Wen and Nakhleh, 2018) implemented in PhyloNet v3.6.5 (Than et al., 2008) was used, as it takes branch lengths and time scale into account for the inference of hybridization and inheritance probabilities. This analysis using the MSNC model was set up for all 20 taxa for 38 nuclear loci and analyzed under default settings. To account for mixing within chains and convergence among chains with reversible jump MCMC (Elworth et al., 2018), a total of 16 chains were run from different seeds for 100 million generations each, with a burn-in preset to 40 million generations and sampling every 10,000 generations. Chains were checked for proper mixing using effective sample size (ESS) > 200 and convergence using the log posterior probability estimated as within-chain average. We ranked the chains according to log posterior probability and discarded chains that differed by more than 10 units of standard deviation of the cumulative set of chains with higher log posterior probability. Phylogeny statistics were obtained from the combined output of the included chains using the SummarizeMCMCResults function in PhyloNet and extracted using Tracer v1.7.1. Owing to the high number of topologies having similar posterior probabilities, we concluded that the most common topology was a better representation of the ‘true’ species phylogeny than the topology happening to have the highest absolute posterior probability. Branch lengths (τ) and coalescent units (θ/2) were translated from units of number of mutations per site to absolute time (Ma) by *post hoc* calibrating the *Hordeum* stem node to 15.3 Ma (Marcussen et al., 2014).

In order to confirm the results from the 38-gene dataset, we replicated the MCMC_SEQ analysis on a 70-gene subset (subsampling every forth gene) of the 275-gene dataset of Marcussen et al. (2014) for 10 species. Analyses were performed with the same settings as before, except the exponential prior on the divergence times of nodes was enabled (-ee) for faster convergence given the higher number of genes (70 vs. 38), and a total of six chains with different seeds were run for 60 million generations each with a burn-in of 20 million generations.

In order to test for additional hybridization that might have become masked by the hybrid origin of the D-genome lineage (see 3. Results) we also analysed the 38-gene dataset without the D-genome lineage species (i.e. A, B, T and outgroups only) as well as D-genome lineage species and outgroups only. For each of the two analyses, a total of six chains with different seeds were run for 60 million generations with a burn-in of 20 million generations.

Finally, we estimated divergence times for the chloroplast phylogeny to clarify its inheritance in hybridization vs. incomplete lineage sorting (Bernhardt et al., 2017; Li et al., 2019, 2015b; Sandve et al., 2015). Given that haploid chloroplast could not be included in the MCMC_SEQ analysis along with the diploid nuclear sequences, we instead dated the chloroplast phylogeny using priors obtained from the posterior in the full-taxon MCMC_SEQ analysis (Fig. 2A and Supplementary Table S4), using concatenated sequence data of *trnT-trnF* and *ndhF-rpl32*. Hence, seven nodes were calibrated: the *Brachypodium distachyon* stem node (root), the *Hordeum vulgare* stem node, the *Agropyron mongolicum* stem node, the *Secale cereale* stem node, the *Taeniatherum caput-medusae* stem node, the *Triticum-Aegilops* crown node and the D-genome crown node (as tmrca of *Ae. caudata* and *Ae. sharonensis*). These were each given an exponential distribution reflecting the haploid coalescence distribution (mean 2.775 Ma) and an offset based on Marcussen et al. (2014): 44.4 Ma, 15.3 Ma, 9.63 Ma, 8.6 Ma, 7.09 Ma, 6.66 Ma, and 3.26 Ma, respectively. The analysis was set up in BEAUti v1.10.4 (strict clock, HKY+G substitution model, Yule tree prior) and run in BEAST v1.10.4 for 10 million generations with 10% burn-in and sampling every 10,000 steps, which was sufficient for proper mixing (ESS of all parameters > 200). The maximum clade credibility tree with Common Ancestor node heights was generated in TreeAnnotator v1.10.4 and visualized in FigTree.

#### 2.4.3 Topology-based inference of phylogenetic network

The SnaQ method implemented in PhyloNetworks (Solís-Lemus et al., 2017) was used to estimate species network topologies from topology frequencies across gene trees for all combinations of species quartets. We first obtained the 38 input gene trees using BEAST v.1.7.4 (Drummond et al., 2012). All alignments were loaded in a single BEAST analysis, with unlinked, strict clock models, linked substitution model GTR+G, and unlinked Yule trees (i.e. not a *BEAST analysis). The MCMC chain was run for 100 million generations with trees sampled every 10,000 generations. The first 20 million generations were discarded from each posterior file as burn-in and the 50% consensus tree was calculated for each of the 38 genes using a custom R script. As recommended by Solís-Lemus et al. (2017), we ran two PhyloNetworks analyses using SnaQ on all 4845 quartets: 10 replicates assuming no hybridization, using the major species tree obtained with MCMC_SEQ as starting tree (see below), *vs*. 20 replicates assuming one hybridization, using the tree with the lowest log pseudolikelihood as starting tree.

#### 2.4.4 SNP-based inference of phylogenetic network using D-statistics

ABBA-BABA tests (or D statistic; Durand et al., 2011; Green et al., 2010; Patterson et al., 2012) were used to test whether a subset of four taxa with phylogeny (((H1,H2),H3),outgroup) contains an excess of single nucleotide polymorphisms (SNPs) indicative of gene flow. Excess of sites with ABBA pattern indicates H2-H3 hybridization, whereas excess of sites with BABA pattern indicates H1-H3 hybridization, but the test cannot identify the direction of gene flow. As backbone tree, we used the major species tree from PhyloNet (Fig. 1B), which has the rooted topology ((A,D),(B,T)), combined with SNPs in the concatenated 38-gene sequence alignment, to test the following four hypotheses: (1) A or D were affected by introgression from B; (2) A or D were affected by introgression from T; (3) B or T were affected by introgression from A; or (4) B or T were affected by introgression from D. In all cases we used *Hordeum vulgare* as outgroup. Iteration over all possible taxon combinations between the four lineages (i.e. the three A taxa, two B taxa, nine D taxa and one T taxon) resulted in 105 tests: 54 tests of hypothesis 1, 27 of hypothesis 2, six of hypothesis 3, and 18 tests of hypothesis 4. ABBA-BABA tests were performed using the *evobiR* package (Blackmon and Adams, 2015) in R cran. Each test was subjected to 1000 jackknife replicates with block size of 2000 bp (~2× length of longest gene) to get an estimate (ŝ) of the standard deviation of D, which was needed to calculate the averaged Z-score and p-value accounting for variance within lineages for each hypothesis.

**Fig. 1.**
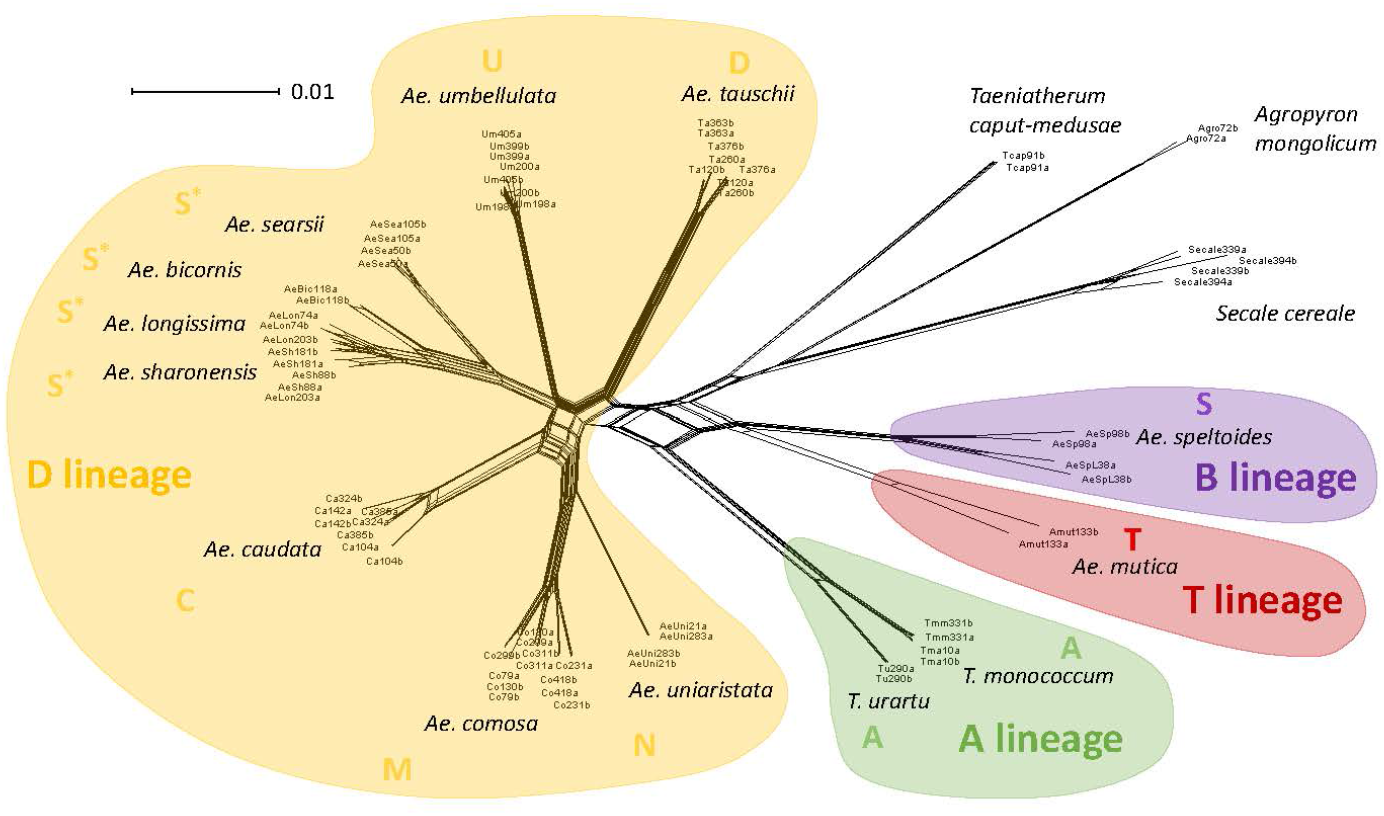
Splits network of diploid Aegilops-Triticum species. Each diploid individual is displayed based on the number of SNP differences among concatenated sequences at 38 nuclear loci. Species names and their genome composition are indicated next to corresponding accessions. S* refers to all variants belonging to Ae. sect. Sitopsis excluding Ae. speltoides (S). Species belonging to the A-, B-, D- or T-genome lineages are colored accordingly.

#### 2.4.5 Estimation of the species major tree with ancestral population sizes

When the species phylogeny is a network, the major species tree is obtained by suppressing the minor hybrid edge (with inheritance probability γ < 0.5) at each reticulation (Solís-Lemus et al., 2017). Assuming complete sampling of lineages, the occurrence of hybridization causes the effective population size to become inflated at nodes ancestral to the hybridization event(s) and as deep as the most recent common ancestor (tmrca) of the parental lineages (species tree dilation; Leaché et al., 2014). Hence, the presence and placement of population size variation among nodes in the phylogeny can be used to generate hypotheses of hybridization. We obtained the major tree by running a phylogenetic analysis allowing for variable population size among branches while enforcing a tree topology (i.e. zero hybridization). The analysis was set up for all taxa for 38 nuclear loci and analyzed in PhyloNet v3.6.5 (Than et al., 2008) using the MCMC_SEQ method (Wen and Nakhleh, 2018) under default settings, except variable population size (-varyps) and maximum number of reticulations set to zero (-mr 0). The MCMC chain was run for 300 million generations, with a burn-in preset to 100 million generations, and sampled every 10,000 generations, which was sufficient for proper mixing (ESS > 200). The results file containing the posterior distribution of trees was converted with a custom GAWK script (https://github.com/aheimsbakk/p2t) to a format compatible with TreeAnnotator v1.10.4 (BEAST v1.10 package; Suchard et al., 2018), through which a maximum credibility tree with common ancestor heights was summarized. The resulting tree and node-specific population size was visualized in FigTree v1.4.3 (http://tree.bio.ed.ac.uk/software/figtree/).

When genes are analysed under the MSC (i.e. under a species tree topology), hybridization typically results in overestimates of population size (species tree dilation) and underestimates of species divergence times (species tree compression) for the affected nodes (Elworth et al., 2018; Leaché et al., 2014; Wen and Nakhleh, 2018). Inclusion or exclusion of hybrids or parental taxa from analyses may therefore result in diagnostic patterns of conflict in population size and node heights among nodes. To explore this, and thereby obtain independent evidence for hybridization in the wheat lineage, we conducted six analyses on each pairwise combination of the four wheat lineages (A, B, D and T). These six analyses (i.e. A/B, A/D, A/T, B/D, B/T and D/T) were run with similar settings as for the all-taxon analysis.

## 3. Results

### 3.1 Genotyping of nuclear markers

Amplicon sequencing of the 48 nuclear loci among the 38 individual samples yielded about 3.9 million of Illumina MiSeq reads that were filtered and trimmed to 1.7 million reads (NCBI XXX). After stringent filtering, it yielded 38 robust loci that were further analyzed. As expected, individual length of sequenced haplotypes ranged from 264 to 535 bp with an average of 361 bp.

### 3.2 Phylogenetic analyses

Global visualization of concatenated sequence data in a splits network recovered the *Aegilops-Triticum* species as distinct from outgroups and comprising four main genetically distinct clusters (i.e. A, B, D and T) whose inter-relationships were unresolved due to a complex, basal split (Fig. 1). Subclusters within the D-genome lineage match taxonomic treatments (i.e. genome U in sect.*Aegilops*, M and N in sect. *Comopyrum*; C in sect. *Cylindropyron*; S* in sect. *Sitopsis* and D in sect. *Vertebrata*), except for the isolated placement of *Ae. speltoides* (S; apart from the other species in sect. *Sitopsis*), and in agreement with downstream phylogenetic analyses.

Summary parameters for the 16 MCMC chains run under the full-parameterized inference of phylogenetic network in MCMC_SEQ are given in Supplementary Table S4. Four chains with highest average posterior log probabilities converged on similar values (–55935 to –55933) and only these topologies were thus further used in downstream analyses. The remaining 12 chains were indeed trapped on lower optima, reaching lower posterior log probabilities (between –55954 and –55945) and being >13.5 units of standard deviation smaller than the cumulative average of ‘best’ chains (Supplementary Table S4). The 95% credible set of topologies for each of the four converging chains comprised between 310 and 614 very similar topologies, reflecting the low node support in parts of the phylogeny (Fig. 2B). All topologies contained the D = A x B reticulation. The same two topologies had the highest incidence in all four chains, overall 4.4% and 3.8% of the total (Supplementary Table S5). The most common network topology, with mean node heights, is shown in Fig. 2A.

**Fig. 2.**
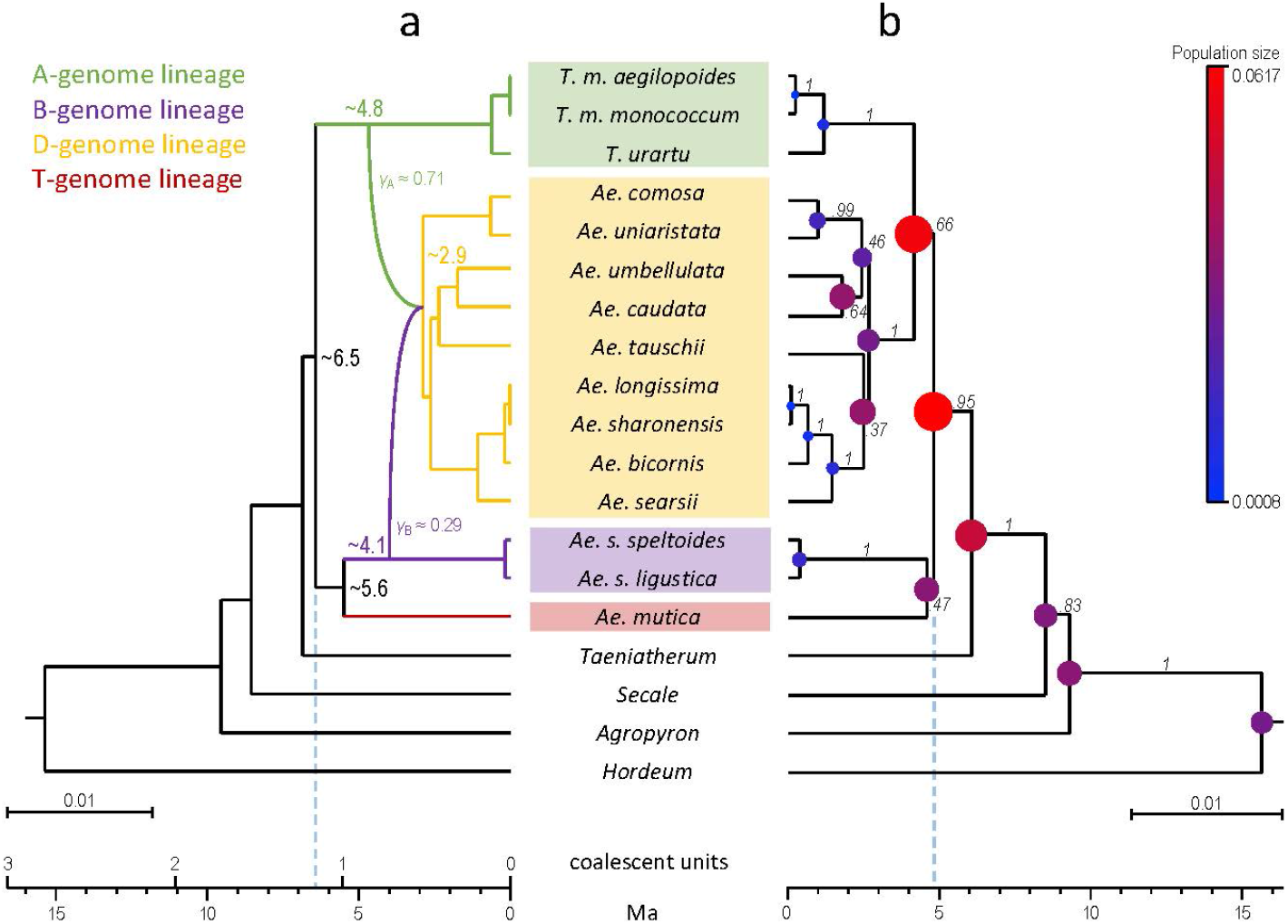
Diploid wheat phylogenies obtained using MCMC_SEQ on a dataset of 38 low-copy nuclear genes. The distant outgroup Brachypodium was pruned from the plots. Scale bars are in units of number of mutations per site. The absolute time scale is interpolated from the Hordeum stem node of 15.3 Ma and obtained ages are shown for nodes of interest. The vertical dotted line projects the wheat crown node age onto the scale axis. A. Phylogenetic network with the highest likelihood, obtained under constant population size and allowing for hybridization. Inferred inheritance probabilities (γA and γB) is indicated for each parent. B. Phylogenetic tree with the highest likelihood, obtained with varying population size among branches and no hybridization. Branch support is indicated as pp. Node balloon size is proportional to population size estimated as units of population mutation rate per site. The young age estimate and high population size estimate for e.g. the wheat crown node reflect tree contraction and tree dilation, respectively.

Inheritance probabilities γ_A_ and γ_B_ supported A as major parent (γ_A_ mean 0.71) and B as minor parent (γ_B_ 0.29), without overlap of 95% credibility intervals (γ_A_ 0.51–0.89 *vs*. γ_B_ 0.11–0.49). Accordingly, the BD node was consistently younger (4.1 Ma; 3.1–5.3 Ma) than the AD node (4.8 Ma; 4.2–5.4 Ma). Average population size (θ) was estimated to 0.0227 (95% credibility interval: 0.0200–0.0254) in units of population mutation rate per site. Average coalescence time, i.e. one coalescent unit, was estimated to 5.6 Ma. All these parameters were very similar across the 10 most common trees (Supplementary Table S5). *Post hoc* analyses on species subsets, i.e. the ‘all taxa except the D-genome lineage species’ and ‘only D-genome lineage species’ subsets (both using *Hordeum* as outgroup), did not reveal additional hybridization (not shown).

Our re-analysis of the data from Marcussen et al. (2014) on a subset of 70 genes using MCMC_SEQ, fully confirmed the herein reported findings in the full-taxon with 38-genes. Four of the six chains converged on the same 95% credible set of 1–2 network topologies, all showing a D = A x B network with A as major parent (γ_A_ mean 0.69–0.74) and a younger BD node than AD node. The remaining two chains failed to mix and had low ESS scores.

The BEAST analysis of the chloroplast, based on priors obtained from the posterior of the full-taxon 38-gene MCMC_SEQ phylogeny (Supplementary Fig. S1), showed a very shallow coalescence of the T-genome lineage within the D-genome lineage, i.e. *Ae. mutica* with *Ae. umbellulata*. Such a significantly more recent coalescence (0.4 Ma; 0.0–1.1 Ma) than their speciation is supportive of introgression. Conversely, the chloroplast for the B-genome lineage coalesced deeply with the rest of wheat species (mean 7.6 Ma; 95% HPD 6.7–8.1 Ma), while the chloroplast of the A- and D-genome lineages coalesced around the time of their speciation (5.4 Ma; 3.9–6.9 Ma).

Topology-based inference of phylogenetic network using SnaQ resulted in a total of 12 network topologies that mostly indicated hybrid origins of single species, but none complying with the D = A x B network inferred by MCMC_SEQ (not shown). SNP-based inference of phylogenetic network using D-statistics (from ABBA-BABA tests) also gave weak and conflicting signals, supporting gene flow between A and T when analyzed with B (hypothesis 3, p = 0.011) but not with D (hypothesis 2, p = 0.657) (Supplementary Table S3).

### 3.3 Estimation of the species major tree and ancestral population sizes

The obtained phylogeny resolved *Brachypodium, Hordeum* (posterior probability; *pp* = 1), *Agropyron* (*pp* = 0.83), *Secale* (*pp* = 1) and *Taeniatherum* (*pp* = 0.95) as successive sisters to an expanded wheat lineage that comprised four main clades (Fig. 2B): the A-genome lineage (with *T. monococcum* and *T. urartu*; *pp* = 1), the B-genome lineage (with two varieties of *Ae. speltoides*; *pp* = 1), the D-genome lineage (with the nine remaining *Aegilops* diploids; *pp* = 1) and the T-genome lineage (with *Ae. mutica*).

The resolution at the base of the wheat lineage was poor, with B and T clustering (*pp* = 0.47) and A and D clustering (*pp* = 0.66). The basal-most branches in the D-genome lineage were short and poorly supported but comprised four main clades: *Ae. tauschii vs. Ae. uniaristata* and *Ae. comosa* (*pp* = 0.99) *vs. Ae. umbellulata* and *Ae. caudata* (*pp* = 0.64) *vs. Ae. searsii, Ae. bicornis, Ae. longissima* and *Ae. sharonensis* (*pp* = 1.0). Most nodes had inferred mean effective population size between 0.0008 (within *T. monococcum*) and 0.0365 (at the base of the D-genome lineage). Three consecutive nodes at the base of the four wheat lineages A, B, D and T presented particularly inflated population size between 0.0481 and 0.0617, indicating possible hybridization between lineages in this part of the major tree.

By iterative exclusion of one of the four wheat genome lineages, the six pairwise analyses of each of the four wheat genome lineages confirmed the hybridization signal by patterns of tree dilation and tree compression (i.e. A/B, A/D, A/T, B/D, B/T, D/T; Supplementary Fig. S2). Comparison of the mean node height and mean population size for each homologous node in the six pairwise-taxon analyses and the all-taxon analysis indeed showed highest variation for both signals for tmrca(A,B,D,T) node (Fig. 3). In particular, analyses comprising D had consistently higher population size (>0.05) for this node than analyses without D (<0.05) and the pairwise analysis of B and D returned the highest population size and lowest age for this node. Such a pattern of tree dilation is consistent with D being a hybrid lineage with A as major parent and B as minor parent.

**Fig. 3.**
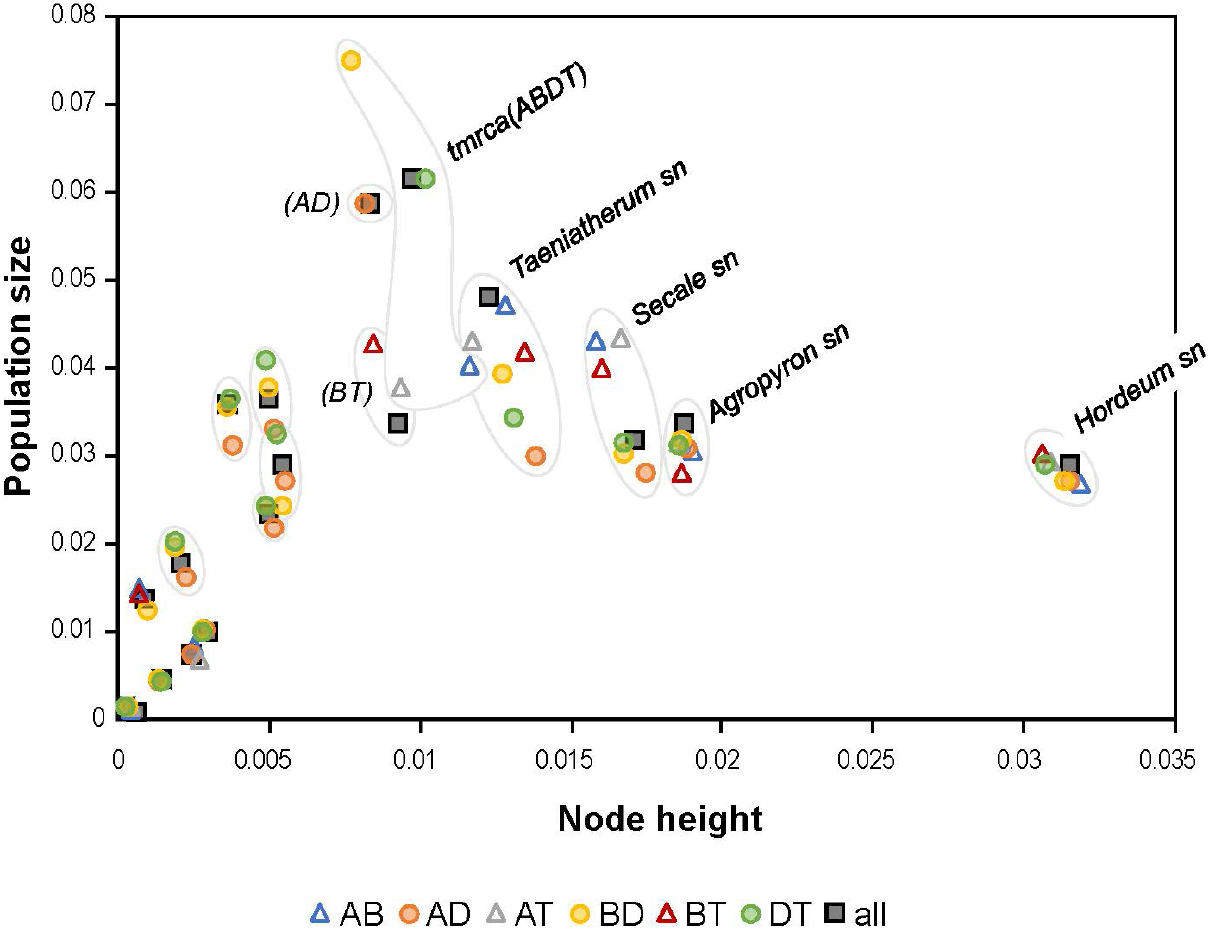
Scatter plot showing dilation of population size and compression of node heights in six pairwise-taxon and one all-taxon phylogenetic analyses. Each analysis was constrained as a tree despite possible hybridization and homologous nodes were fenced and named (for those preceding the D = A x B hybridization). Datapoints for the four analyses comprising D (i.e. all, AD, BD, DT), and thus containing hybridization, are indicated with filled symbols. Population size and node heights are displayed as units of population mutation rate per site and as units of number of mutations per site, respectively. Abbreviations: sn = stem node.

## 4. Discussion

We here present the first comprehensively sampled, dated phylogeny for wheat and relatives, based on full-parameterized coalescent analyses on sequences of 38 low-copy loci for 13 diploid *Aegilops-Triticum* species. We recovered this wheat lineage as monophyletic, with *Taeniatherum* as its closest sister, and comprising four main diploid sublineages interconnected by hybridization in a phylogenetic network, settling a long-standing debate on the delimitation of the wheat lineage (reviewed in Feldman and Levy, 2015). We propose to refer to these sublineages as the A-genome lineage (i.e. *Triticum* diploids), B-genome lineage (i.e. *Ae. speltoides*), the T-genome lineage (i.e. *Ae. mutica*), and the D-genome lineage (nine diploid *Aegilops* species), expanding on the simplified terminology proposed by Marcussen et al. (2014). Our phylogeny agrees well with predictions based on morphological character states indicating that *Ae. mutica, Ae. speltoides, T. monococcum* and *T. urartu* mostly exhibit ancestral traits in various combinations (Eig, 1929a, 1929b). Contrastingly, species in the hybrid D-genome lineage exhibit advanced traits such as subtelocentric chromosomes (present in C/M/N/U) derived from more metacentric chromosomes (present in T/A/S/D; Feldman and Levy, 2015). This D-genome lineage was found here to unite diploid *Aegilops* species with divergent karyotypes and differential pairing at meiosis (Kihara, 1954), that were previously placed in five different sections of the genus (i.e. *Ae*. sect. *Sitopsis* with S* genome, sect. *Aegilops* with U, sect. *Cylindropyron* with C, sect. *Comopyrum* with M and N, and sect. *Vertebrata* with D, summarized in Feldman and Levy, 2015). The sister relationship between *Ae. mutica* and *Ae. speltoides* is also supported by a number of characters, such as allogamy, meiotic pairing of their hybrid, a suppressor of the Pairing homeologous gene (Ph1) or the presence of B-chromosomes (reviewed in Ohta, 1991; Feldman and Levy, 2015).

Hybridization set aside, our species phylogeny delimitates the main lineages differently than previous studies that employed fewer genes (the 6-gene phylogeny in Marcussen et al., 2014; Petersen et al., 2006), but generally agrees with obtained cpDNA phylogenies (e.g. Bernhardt et al., 2017; Li et al., 2015a, 2015b; Sandve et al., 2015). Discrepancies may be attributable to coalescence stochasticity or chloroplast introgression as also here detected from *Ae. umbellulata* into *Ae. mutica* (Supplementary Fig. S1). Although the wheat lineage itself is recovered as monophyletic, neither of its two nested genera, *Triticum* and *Aegilops* (including *Amblyopyrum*), is clearly delimited. They are interwoven by chloroplast introgression (Supplementary Fig. S1), homoploid hybridization (Fig. 2) and allopolyploidy (Kilian et al., 2011), supporting their proposed unification in an expanded *Triticum* genus (e.g. Feldman and Levy, 2015; Stebbins, 1956; but see van Slageren, 1994).

### 4.1 The hybrid origin of the D-genome lineage

The two main lines of evidence presented here (Figs 2 and. 3) both support that the ancestor of the D-genome lineage was a homoploid hybrid between the A-genome lineage and the B-genome lineage. This confirms previous findings based on fewer taxa with 275 gene trees (Marcussen et al., 2014) or transposable elements (El Baidouri et al., 2017). While Marcussen et al. (2014) concluded with almost equal parental contributions to the hybrid based on tree topology ratios and similar ages of splits, our finding herein as well as the re-analysis of a 70-gene subset of the data of Marcussen et al. (Supplementary Fig. S3) indicate that gene flow from the B-genome lineage into D was smaller (γ_B_ mean: 0.29; 95% CI 0.11–0.49) and ~0.7 Ma more recent. Such refinement highlights the superior ability of the full-parameterized approach based on explicit time scales to distinguish between scenarios of small-scale but recent introgression *vs*. large-scale but ancient introgression that topology-based and SNP-based methods otherwise confound. It is noteworthy that neither the topology method (SNaQ; Solís-Lemus et al., 2017) nor the SNP ratio method (ABBA-BABA tests; Durand et al., 2011; Green et al., 2010) was able to detect the hybridization event for this 38-gene dataset, although readily identifiable by the MCMC_SEQ method (Figs. 2 and 3).

Our dated phylogeny suggests an evolutionary scenario involving an initial speciation event some 6.5 Ma ago that gave rise to the A-genome lineage and another lineage that rapidly yielded B- and T-genome lineages (5.6 Ma ago). A ‘proto-D-genome’ lineage diverged from the A-genome lineage about 4.8 Ma ago and was soon after (~4.1 Ma) massively affected by gene flow from the B-genome lineage, yielding the modern admixed D-genome lineage with A as major parent (γ_A_ ≈ 0.71) and B as minor parent (γ_B_ ≈ 0.29). This reticulation event was followed by substantial radiation of this D-genome lineage from ~2.9 Ma onwards. Whether this reticulation corresponded to massive introgression or hybrid speciation remains arguable. Given that introgression implies modification of an existing lineage by retention of usually a few loci from another species, the massive proportion of genes transferred from the minor parent B into the D-genome lineage, contributing possibly one third of its genome, rather seems consistent with origination of a new lineage by hybrid speciation (Folk et al., 2018; Mallet, 2007). Recombinational speciation is indeed likely to yield unequal genetic contributions from the parental species, as cytonuclear interactions or genome dominance can bias the long-term maintenance of maternal *vs*. paternal loci (Freeling et al., 2015). Such processes have been reported to play a role in the evolution of wheat genomes (Feldman and Levy, 2015; Li et al., 2019; Senerchia et al., 2015) and should be further investigated regarding the D-genome lineage.

### 4.2 Is there any evidence of additional hybridization in the origin of the D-genome lineage?

Following the initial evidence of an hybrid origin of the D-genome lineage (Marcussen et al., 2014), subsequent studies have suggested “more complex” hybridization scenarios, however without presenting formal analyses in support of a different scenario (El Baidouri et al., 2017; Li et al., 2015a, 2015b). Our new data provide no evidence for additional hybridization other than the recent chloroplast introgression from *Ae. umbellulata* into *Ae. mutica* (Supplementary Fig. S1). None of the remaining nodes displayed a distinctly inflated population size that would be suggestive of tree dilation caused by hybridization (Leaché et al., 2014) and none of the MCMC_SEQ analyses on data subsets inferred additional hybridization events. Of course, additional hybridization events cannot be strictly ruled out but, to remain undetected by our study, must have been minor and involve recently diverged parents or smaller γ than in the D = A x B event. With an average time to coalescence (i.e. 5.5 Ma) approaching the age of the entire wheat lineage (6.5 Ma), additional hybridization events at the origin of the D-genome lineage certainly remain a challenge to detect (Elworth et al., 2018). Under such conditions SNP-based or topology-based methods are of little use because the vast majority of genes (>70%; Marcussen et al. 2014) coalesce more deeply than tmrca and confer noise or, even worse, bias. Given the heavy computational load of available full-parameterized approaches for the many genes required for proper resolution, analysis of introgression in such species systems may have to rely on phylogenetic approaches yet to be developed.

### 4.3 Diversification of the hybrid D-genome lineage

With nine diploid species, the hybrid D-genome lineage outnumbers the three other wheat lineages which together comprise a total of four diploid species. This D-genome lineage has expanded beyond the overlapping ranges of the parental lineages in the Fertile Crescent and greatly diversified across Mediterranean landscapes (Kilian et al., 2011). The predominantly selfing reproductive regime of D may have promoted speciation (Grundt et al., 2006; Wright et al., 2013), although this lineage has noticeably produced more species than the equally selfing A-genome lineage. Diversification of the D-genome lineage coincides with the onset of Pleistocene climatic oscillations that may have promoted speciation (Levin, 2002). It was accompanied by considerable changes in genome size (Eilam et al., 2007) and organization (Badaeva et al., 1996) that were consistent with either upsizing of metacentric chromosomes (e.g. among S*-genome species) or genome downsizing associated with chromosomal asymmetry in species such as *Ae. umbellulata*. Such karyotype evolution was predicted to support adaptive radiation (Stebbins, 1971), but neither underlying mechanisms nor significance for diversification are yet understood. Given that this radiation was preceded by genetic admixture, the underpinning driver may be associated directly with hybridization (Marques et al., 2019). Future work will possibly address to what extent it promoted recombination of adaptive variation from parental lineages (Keller et al., 2013) and/or fostered the origin of innovative traits (Abbott et al., 2013).

## Acknowledgments

This work was funded by the Swiss National Science Foundation (31003A-153388). Data analyzed in this paper were generated at the Genetic Diversity Centre, ETH Zurich. We thank A. Heimsbakk for providing the GAWK script as well as S. Grünig and reviewers for useful feedbacks on the manuscript.

## Author contributions

SH, FF and CP designed the study. SH collected data; SH and TM analyzed data; SH, TM, FF and CP wrote the paper. All authors approved the final version of the manuscript.

## Supplementary Material (on-line only)

**Fig. S1:** Chloroplast introgression from *Ae. umbellulata* into *Ae. mutica*

**Fig. S2:** Full-parameterized species trees for pairs of the genome-lineages A, B, D, T

**Table S1:** List of accessions of *Aegilops-Triticum* species and outgroups used in this study

**Table S2:** List of the 48 genotyped low-copy loci

**Table S3:** Summary statistics for all-taxon iterations of ABBA-BABA tests

**Table S4:** Statistics of 16 MCMC_SEQ runs from different seeds

**Table S5:** Statistics for the 10 most common topologies for the four best chains combined (separate file)

**Script S1:** R script for Illumina read filtering and cleaning (separate file)

**Figure S1.**
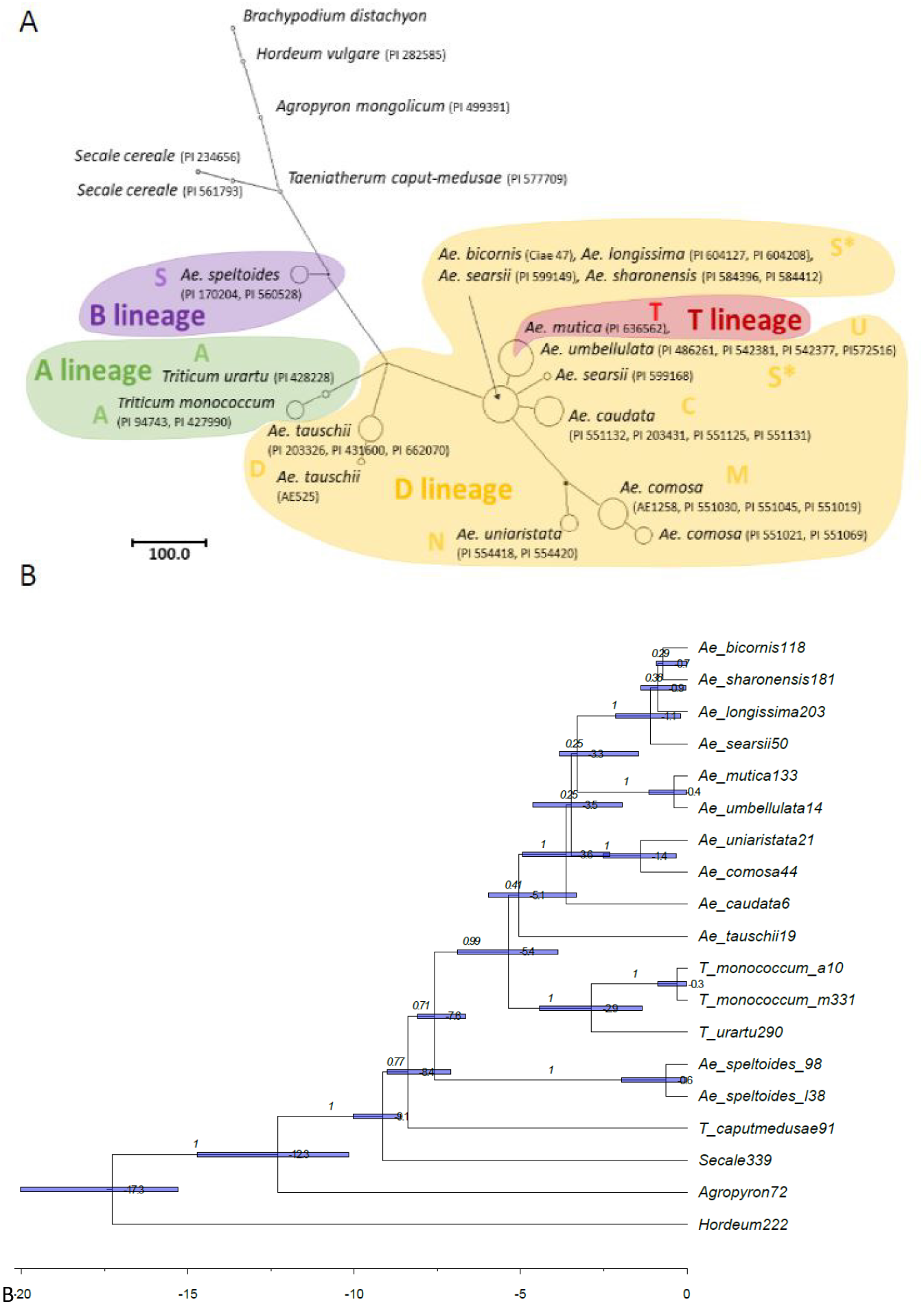
Chloroplast introgression from *Ae. umbellulata* and *Ae. mutica*. **(A)** Haplotype network of two concatenated plastid loci (*trnT-trnF* and *ndhF-rpl32*). Plastid loci were PCR amplified and Sanger sequenced according to Meimberg *et al*. (2009) for each sample (NCBI XXX to XXX) and concatenated into 1760 bp of plastid haplotype. For haplotype network, species names and their genome composition are indicated next to corresponding accessions. S* refers to all variants belonging to *Ae*. sect. *Sitopsis* excluding *Ae. speltoides* (S). Species belonging to the A-, B-, D- or T-genome lineages are colored accordingly. **(B)** Maximum clade credibility BEAST phylogeny of two concatenated plastid loci (*trnT-trnF, ndhF-rpl32*). Calibrations were informed from the most common network topology (Fig. 2A). Node labels refer to average node height, node bars refer to node height 95% PHD, and branch labels refer to branch support as posterior probability. The remote outgroup *Brachypodium* was trimmed from the plot.

**Figure S2 (below).**
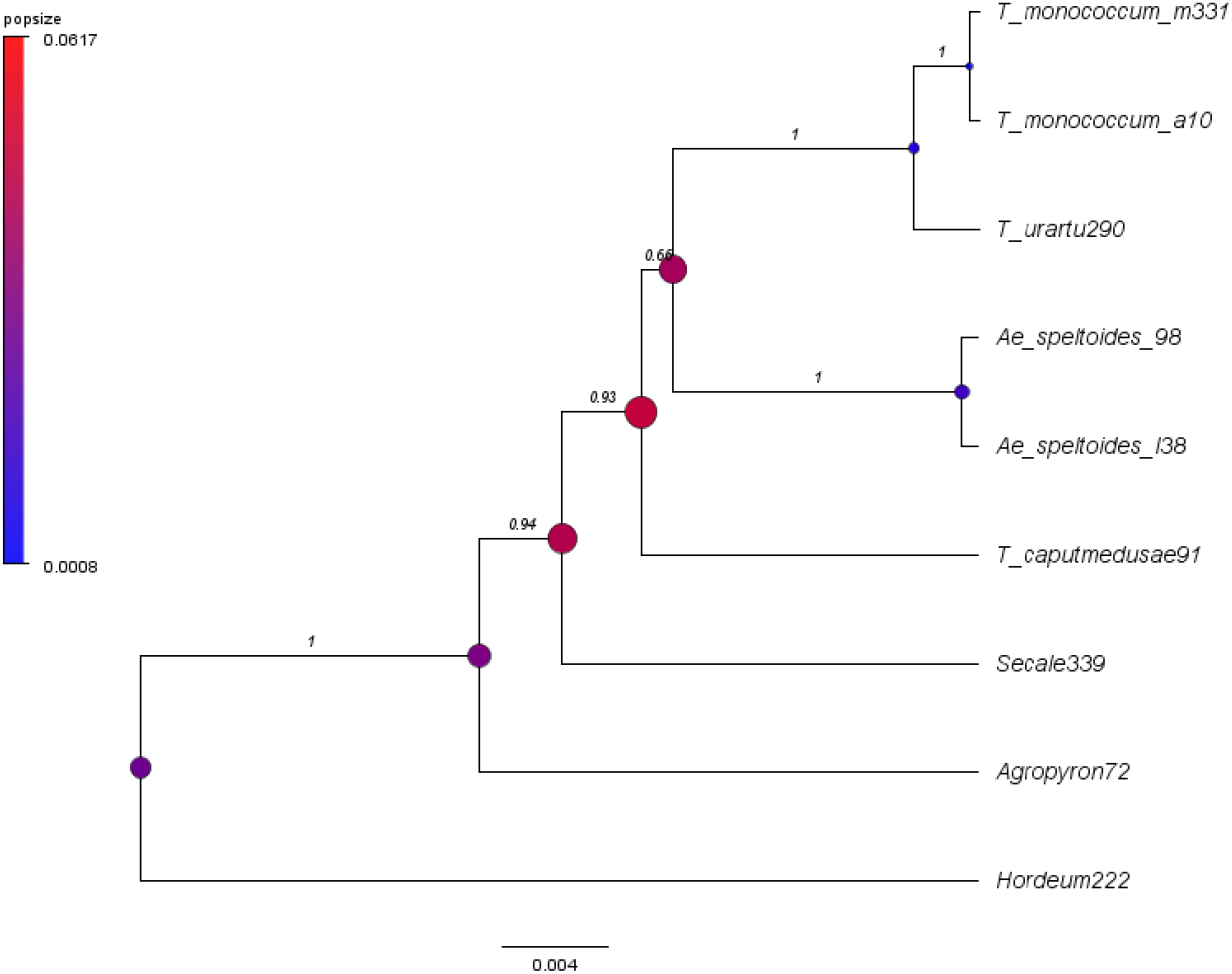
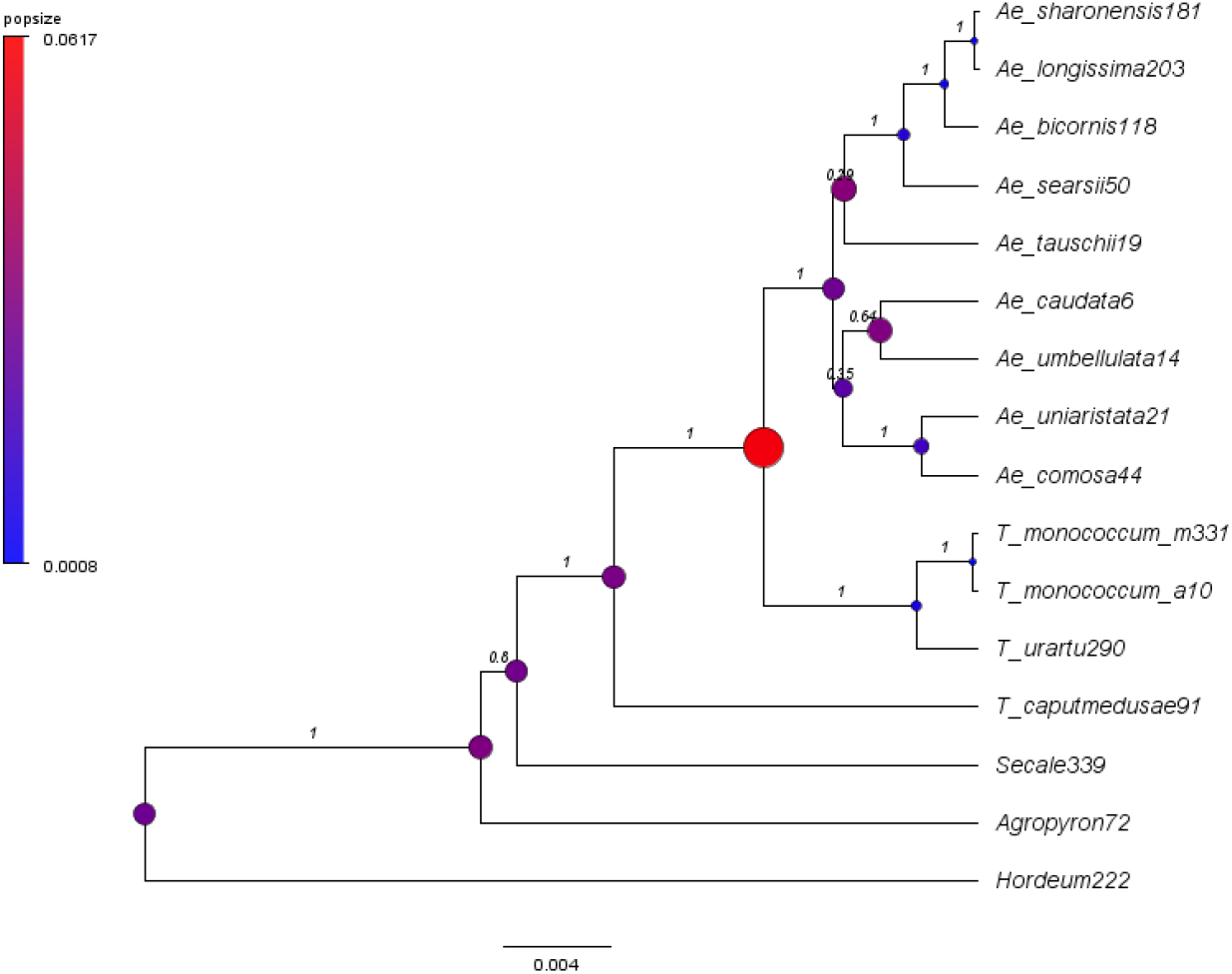
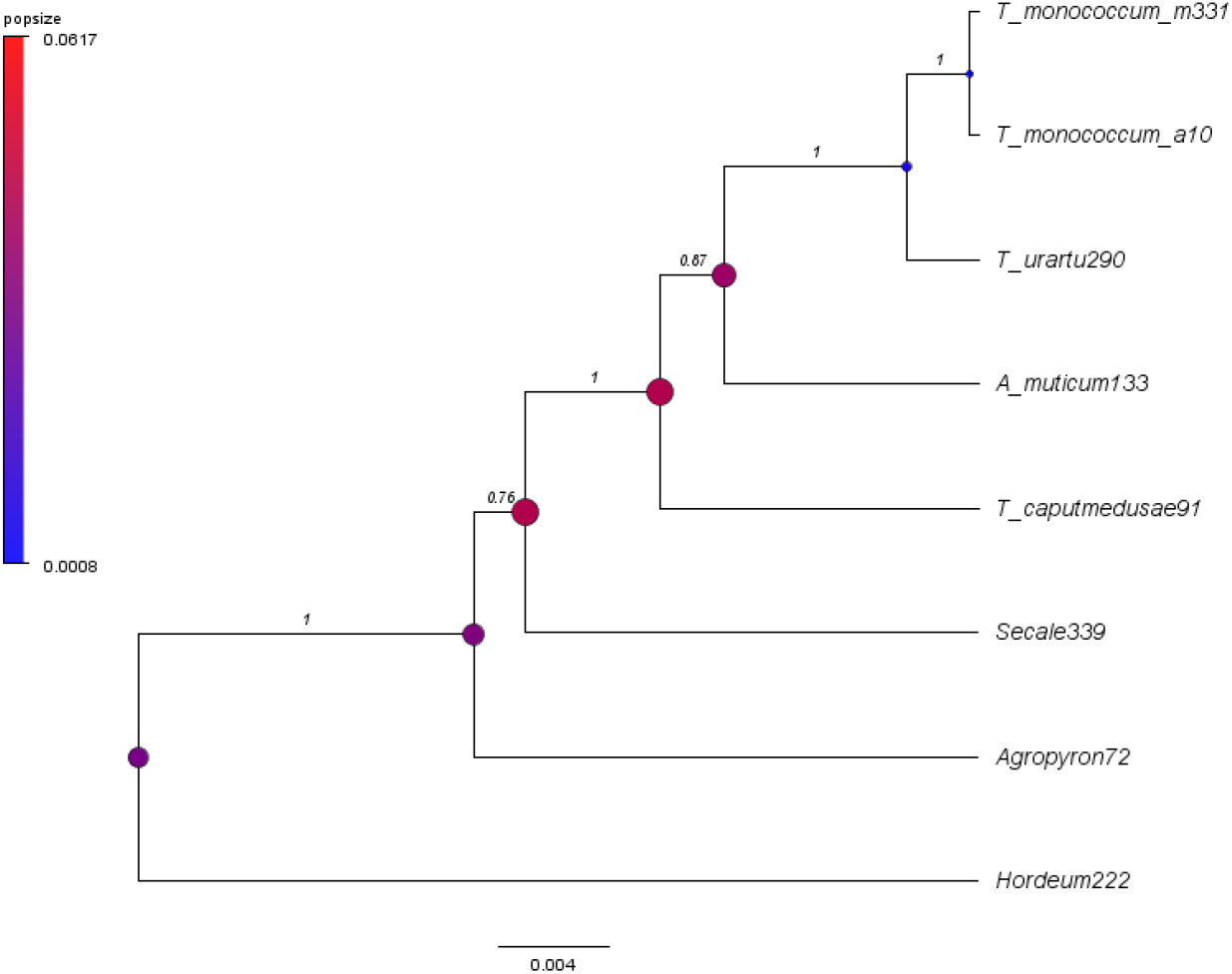
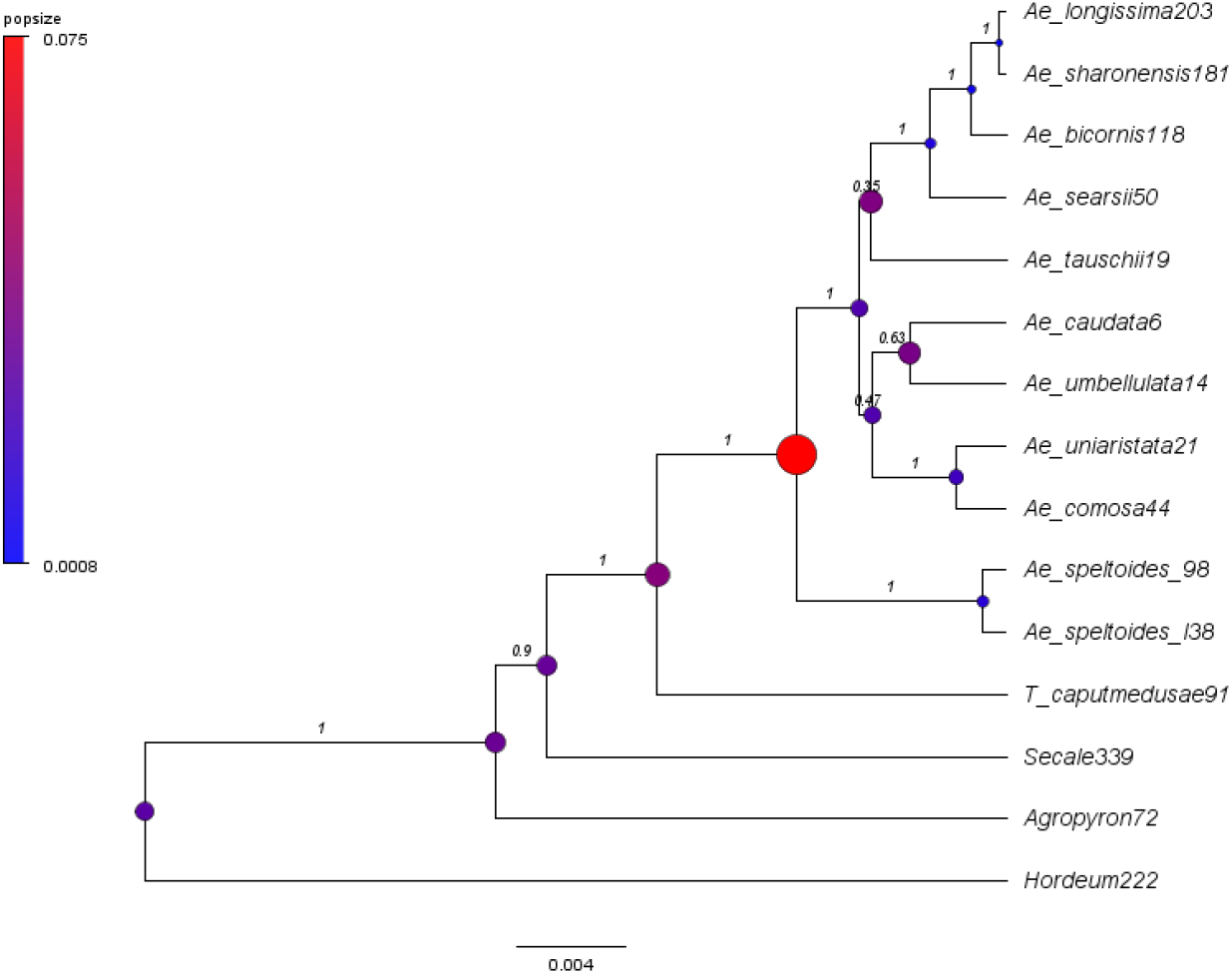
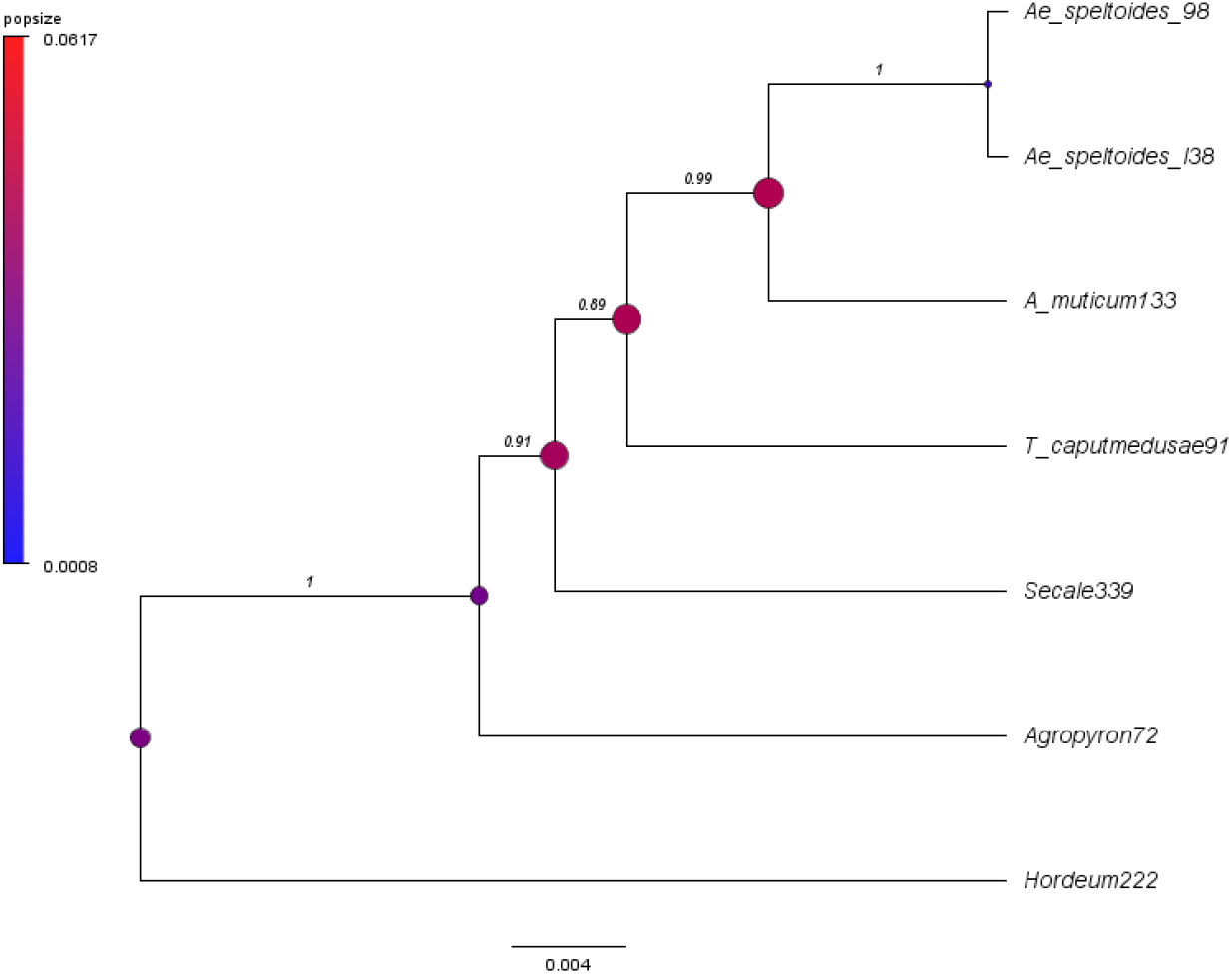
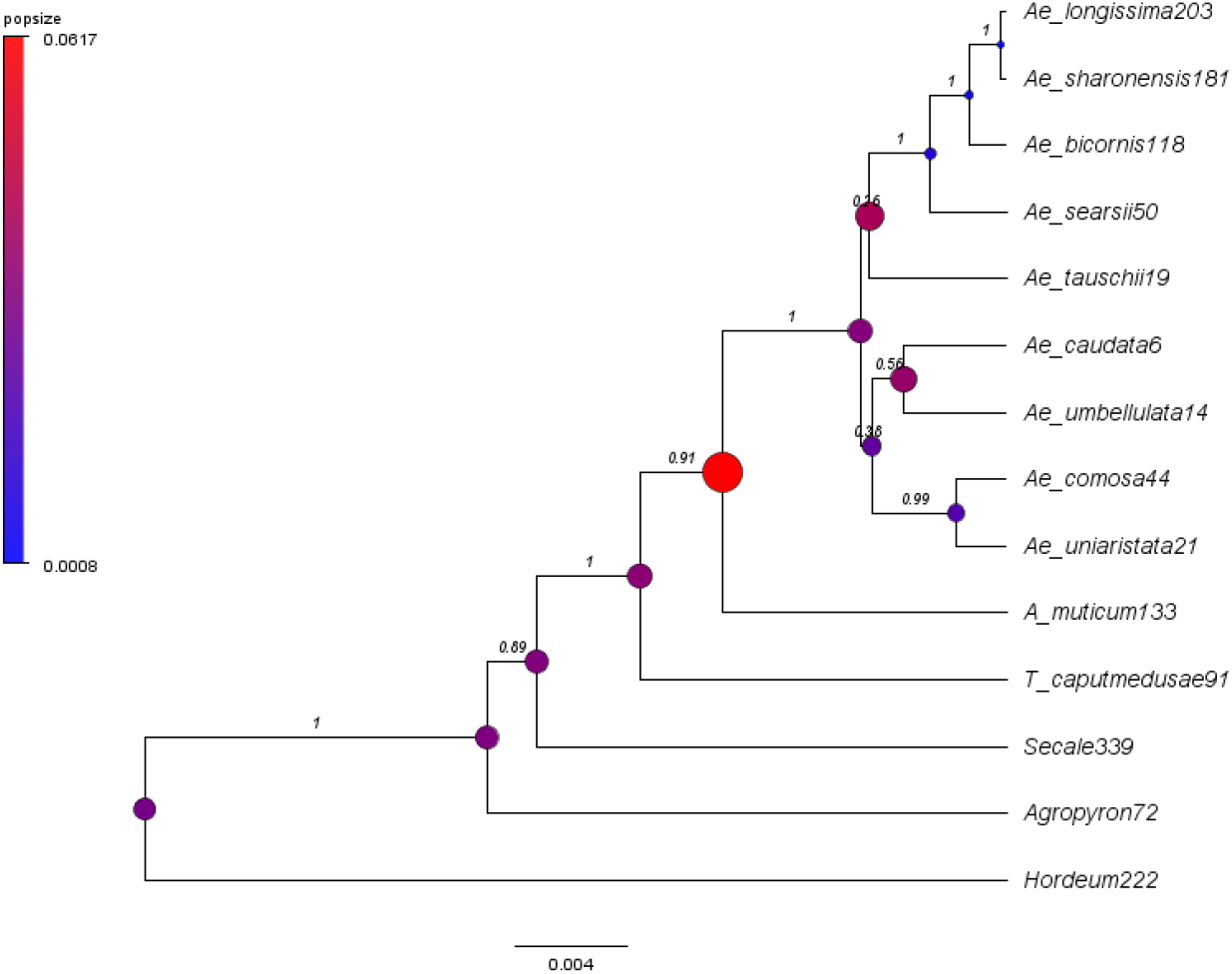
Full-parameterized species trees for the six pairwise analyses of the four genome-lineages A, B, D and T. **(A)** Pairwise analysis of A & B. **(B)** Pairwise analysis of A & D. **(C)** Pairwise analysis of A & T. **(D)** Pairwise analysis of B & D. **(E)** Pairwise analysis of B & T. **(F)** Pairwise analysis of D & T. Trees were inferred from 38 nuclear loci under MCMC_SEQ in PhyloNet, a tree constraint and varying effective population size. Estimated population size and posterior probability are shown for each tree, as node balloons and branch labels, respectively. The remote outgroup *Brachypodium* was trimmed from each plot. The all-taxon tree is shown in Fig. 2B.

**Figure S3.**
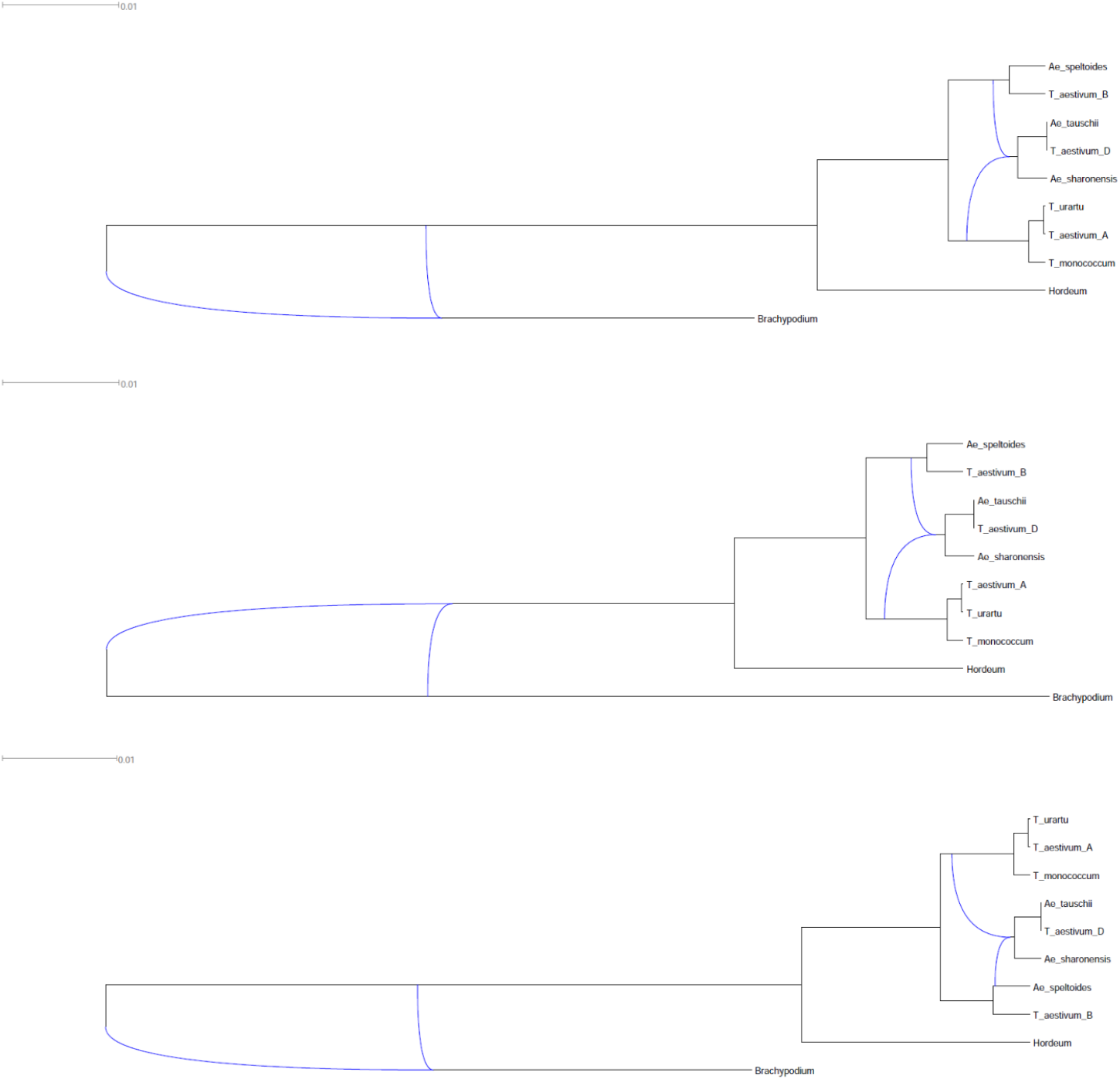
All three network topologies obtained by MCMC_SEQ analysis of a 70-gene subset of the 275-gene dataset of Marcussen *et al*. (2014). The two first topologies, differing only in the placement of an inferred reticulation involving the outgroup, occurred at high frequency (94.0–99.5%) in all the four MCMC chains, while the third occurred at low frequency (2.3%) in one chain only.

**Table. S1.** List of the 38 accessions of the diploid *Aegilops-Triticum* species and outgroups used in this study. The genome composition of species were reported following van Slageren (1994) and Barkworth & von Bothmer (2009). Accessions were provided by USDA ARS-GRIN (PI; CIae) or IPK-Gatersleben (AE) genebanks

| Species | Genome | Accessions |
| --- | --- | --- |
| *Ae. caudata* (= *Ae. markgrafii*) | C | PI 551132, PI 203431, PI 551125, PI 551131 |
| *Ae. comosa* | M | AE 1258, PI 551030, PI 551021, PI 551019, PI 51045, PI 551069 |
| *Ae. tauschii* | D | AE 525, PI 220326, PI 431600, PI 662070 |
| *Ae. umbellulata* | U | PI 486261, PI 542381, PI 542377, PI 573516 |
| *Ae. uniaristata* | N | PI 554418, PI 554420 |
| *Ae. bicornis* | S<sup>b</sup>=S* | Clae 47 |
| *Ae. sharonensis* | S<sup>sh</sup>=S* | PI 584396, PI 584412 |
| *Ae. searsii* | S<sup>s</sup>=S* | PI 599168, PI 599149 |
| *Ae. longissima* | S<sup>l</sup>=S* | PI 604127, PI 604108 |
| *Ae. speltoides* var. *speltoides* | S=B | PI 170204 |
| *Ae. speltoides* var. *lingustica* | S=B | PI 560528 |
| *Ae. mutica* (= *Amblyopyrum muticum*) | T | PI 636562 |
| *T. monococcum* spp. *monococcum* | A<sup>m</sup> | PI 94743 |
| *T. monococcum* spp. *aegilopoides* | A<sup>m</sup> | PI 427990 |
| *T. urartu* | A<sup>u</sup> | PI 428228 |
| *Agropyron mongolicum* | P | PI 499391 |
| *Taeniatherum caput-medusae* | Ta | PI 577709 |
| *Secale cereale* | R | PI 234656, PI 561793 |
| *Hordeum vulgare* | I | PI 282585 |

**Table S2 List of the 48 genotyped low-copy loci (Electronic material)**. Names of loci as annotated in *Triticum aestivum* reference genome, with description and name of corresponding loci in *Brachypodium distachyon* and *Aegilops tauschii*. Reverse and forward primers for amplicon sequencing are presented with corresponding melting temperatures (°C), expected length of targeted sequences (bp) and results of the post-processing quality filtering steps.

**Table S3.** Summary statistics for all-taxon iterations of ABBA-BABA tests. N quartet combinations of A, B, D or T wheat genome lineages with *Hordeum vulgare* as outgroup tested four hybridization hypotheses in the following order: hybridization between B and either A or D, hybridization between T and either A or D, hybridization between A and either B or T, hybridization between D and either B or T.

| Hybridization hypotheses<br>(((H1,H2),H3),O) | N | Mean length<br>(bp) | ABBA sites | BABA sites | % ABBA | mean D | stdev D | Z-score | p-value | Interpretation |
| --- | --- | --- | --- | --- | --- | --- | --- | --- | --- | --- |
| 1:(((A,D),B),*Hordeum*) | 54 | 11'496.3 | 44.24 | 43.78 | 0.50 | 0.004 | 0.115 | 0.090 | 0.464 | No hybridization |
| 2:(((A,D),T),*Hordeum*) | 27 | 11'155.0 | 39.48 | 41.63 | 0.48 | -0.031 | 0.107 | -0.404 | 0.343 | No hybridization |
| 3:(((B,T),D),*Hordeum*) | 18 | 11'150.7 | 42.67 | 35.72 | 0.55 | 0.093 | 0.059 | 1.509 | 0.066 | No hybridization |
| 4:(((B,T),A),*Hordeum*) | 6 | 11'199.5 | 43.00 | 33.00 | 0.57 | 0.131 | 0.054 | 2.290 | 0.011 | Gene flow between A and T |

**Table S4:**
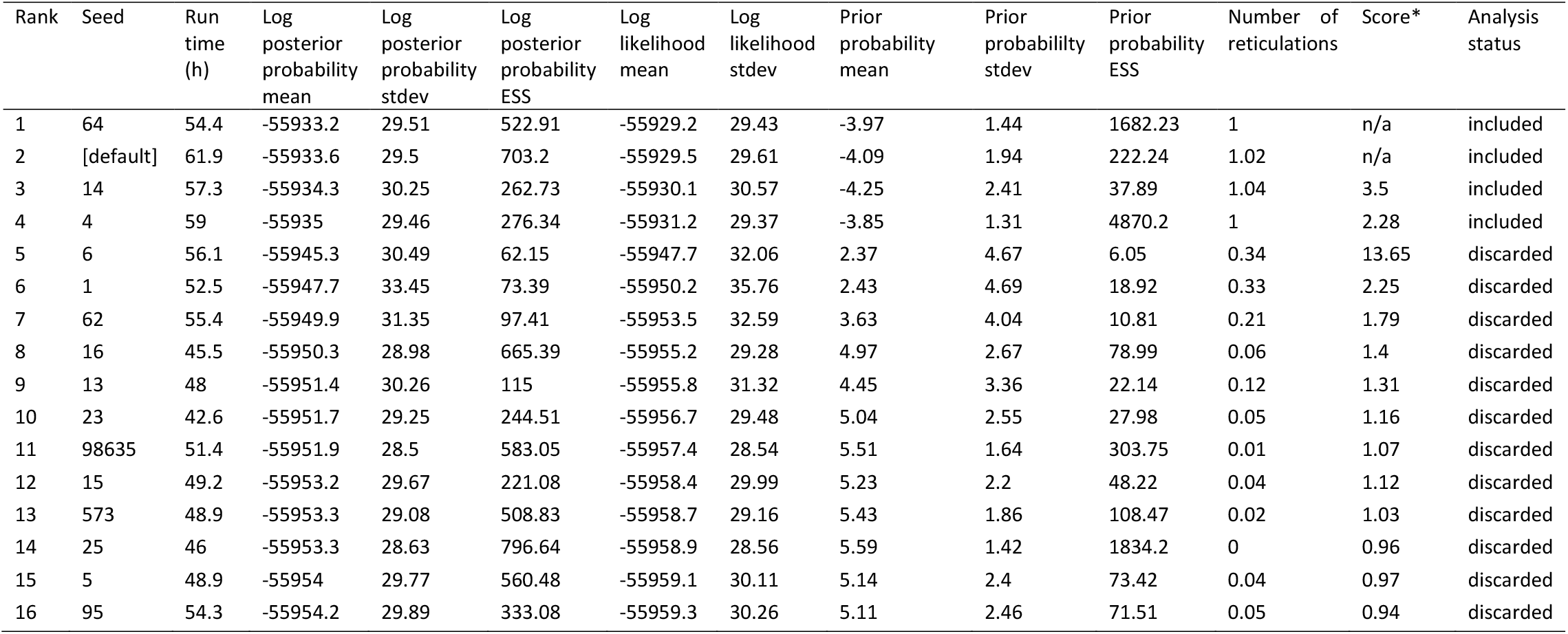
Statistics of 16 MCMC_SEQ runs from different seeds. Chains are ranked by their log posterior probability means after removal of burn-in, from highest to lowest. Only the four best chains were used for phylogeny inference (Supporting Information Table S5).

